# Tropical arboreal ants form dominance hierarchies over nesting resources in agroecosystems

**DOI:** 10.1101/442632

**Authors:** Senay Yibarek, Stacy M. Philpott

## Abstract

Interspecific dominance hierarchies have been widely reported across animal systems. While some dominant individuals (winners) get to monopolize resources, during dyadic interactions, they can increase their relative fitness as compared to subdominant individuals (losers). In some ant species, dominance hierarchies have been used to explain species coexistence and community structure. However, it remains unclear whether or in what contexts dominance hierarchies occur in tropical ant communities. Furthermore, it can be challenging to infer and quantify reliable dominance hierarchies from observed interactions. This study seeks to examine whether arboreal twig-nesting ants competing for nesting resources in a Mexican coffee agricultural ecosystem are arranged in a dominance hierarchy. Using network analysis, we quantified interactions between ten species by measuring the uncertainty and steepness in the dominance hierarchy. We also assessed the orderliness of the hierarchy by considering species interactions at the network level. Based on the Elo-ranking method, we found that the twig-nesting ant species *Myrmelachista mexicana* ranked highest in the ranking, while *Pseudomyrmex ejectus* was ranked as the lowest in the hierarchy. We quantified the uncertainty in the dominance hierarchy and found that the hierarchy was intermediate in its steepness, suggesting that the probability of higher ranked individuals winning contests against lower ranked individuals was fairly high. Motif analysis and significant excess of triads further revealed that the species networks were largely transitive. This study highlights that some tropical arboreal ant communities self-organize into dominance hierarchies.

## Introduction

A long-standing goal in ecology has been to determine the underlying mechanisms that give rise to species coexistence in local communities, especially in assemblages with multiple competing species (MacArthur 1958; Hutchinson 1959). Numerous mechanisms have been proposed for maintaining species coexistence (Wright 2002; Silvertown 2004). Interspecific competitive trade-offs, whereby the superiority of a particular species in an environment or biotic condition is balanced by the inferiority of other species, can lead to segregation among species (Tilman 1994; Levine et al. 2004). These interspecific interactions are thought to lead to the long-term stable coexistence of ecologically similar species (Levins 1979; Holt et al. 1994; Chesson 2000; Bever 2003; Rudolf and Antonovics 2005), and may also be characterized by dominance hierarchies. Dominance hierarchies have been widely observed in a wide range of taxa, from vertebrates to invertebrates (Chase and Seitz 2011). Species can be ranked into dominant species (i.e. higher-ranked individuals) or subordinate species (i.e. lower-ranked individuals) on the basis of aggression or ritual displays (Drews 1993). Interspecific dominance hierarchies have been used to understand patterns of local species coexistence in ecological communities with as consequence that higher ranked individuals monopolize resources resulting in fitness benefits (Morse 1974; Schoener 1983).

In ant communities, dominance hierarchies have been used to examine interspecific tradeoffs to explain species coexistence patterns (Stuble et al. 2013). These trade-offs include the discovery-dominance trade-off, the discovery-thermal tolerance tradeoff, and the discovery-colonization trade-off (Cerdá et al. 1998*a*; Stanton et al. 2002; Lebrun and Feener 2007; Stuble et al. 2013). In addition to testing interspecific trade-offs, dominance hierarchies have been used to understand the role of dominant species in structuring local communities and composition, such as partitioning dominant and subdominant species within guilds (Baccaro et al. 2010; Arnan et al. 2012). Dominant ant species play an important role in the structuring of local communities. For example, *Formica* species dominating a boreal ecosystem divert resources away from subdominant competitors (Savolainen and Vepsäläinen 1988). In Mediterranean ecosystems, subdominant species forage at nearly lethal environmental conditions while dominant species reduce mortality risk by foraging at more favorably temperatures (Cerdá et al. 1998*c*). In tropical ecosystems, competing arboreal ants can be structured into a dominance hierarchy with higher ranked ant species having greater access to nesting sites and extrafloral nectaries. However, levels of uncertainty associated with outcomes of interspecific interactions are often not quantified (Stuble et al. 2017). Furthermore, its remains unclear how arboreal ants or tropical ants are structured at higher-order interactions, such as when interspecific interactions are viewed as a network.

In this study, we examine dominance hierarchies for a community of arboreal twig-nesting ants in a coffee agroecosystem. Both arboreal and ground-dwelling twig-nesting ants in coffee agroecosystems are nest-site limited in terms of number (Philpott and Foster 2005*a*), diversity (Armbrecht et al. 2004; Gillette et al. 2015), and sizes (Jiménez-Soto and Philpott 2015) of nesting resources. Many studies focused on interspecific dominance hierarchies lack a clear and consistent definition of dominance. For twig-nesting ants, nest takeovers are common and nest sites are often limiting, thus dominance in this system is defined as competition for nest sites (Brian 1952), and in at least one case has been experimentally demonstrated (Palmer et al. 2000a). Although dominance studies use various methods to rank species, they don’t typically account for uncertainty in rankings, except for a few cases (Adler et al. 2007; Stuble et al. 2013). This present study aims to develop dominance hierarchies for twig-nesting ants living in hollow twigs in a Mexican coffee agricultural ecosystems due to competition for nest resources. We adopt methods from network analysis to infer dominance hierarchies from competitive interactions by estimating uncertainties and steepness in rankings (Pinter-Wollman et al. 2014; Shizuka and McDonald 2015; Sánchez-Tójar et al. 2017). Furthermore, by viewing twig-nesting ants as a network we estimate the orderliness of the hierarchy within the community. Specifically, we tested the hypothesis that tropical, arboreal twig-nesting ants form a clear, dominance hierarchy for nesting sites in controlled environments.

## Methods

### Study Site and System

We conducted fieldwork at Finca Irlanda (15°20’ N, 90°20’ W), a 300 ha, privately owned shaded coffee farm in the Soconusco region of Chiapas, Mexico with ~250 shade trees per ha. The farm is located between 900-1100 m a.s.l. Between 2006-2011, the field site received an average rainfall of 5726 mm per year with most rain falling during the rainy season between May and October. The farm hosts ~50 species of shade trees that provide between 30-75% canopy cover to the coffee bushes below. The farm has two distinct management areas -- one that is a traditional polyculture and the other that is a mixture of commercial polyculture coffee and shade monoculture coffee according to the classification system of (Moguel and Toledo 1999).

The arboreal twig-nesting ant community in coffee agroecosystems in Mexico is diverse. There are ~40 species of arboreal twig-nesting ants at the study site including *Brachymyrmex* (3 species), *Camponotus* (8), *Cephalotes* (2), *Crematogaster* (5), *Dolichoderus* (2), *Myrmelachista* (3), *Nesomyrmex* (2), *Procryptocerus* (1), *Pseudomyrmex* (11), and *Technomyrmex* (1) (Philpott and Foster 2005*a*; Livingston and Philpott 2010).

### ‘Real-estate’ experiments

We examined the relative competitive ability of twig-nesting ants by constructing dominance hierarchies based on ‘real estate’ experiments conducted in the lab. We collected ants during systematic field surveys in 2007, 2009, 2011, and 2012 in the two different areas of the farm, and then used ants in lab experiments. We first removed ants from individual twigs, and then placed ants (workers, alates, and brood) from two different species (one twig per species) into sealed plastic tubs with one empty artificial nest. The artificial nest, or ‘real estate’, consisted of a bamboo twig, 120 mm long with a 3-4 mm opening. After 24 hours, we opened the bamboo twigs to note which species had colonized the twig. All ants collected were used in ‘real estate’ trials within two days of collection, or were discarded.

We conducted trials between pairs of the ten most common ant species encountered during surveys: *Camponotus abditus*, *Camponotus* (*Colobopsis*) sp. 1, *Myrmelachista mexicana*, *Nesomyrmex echinatinodis*, *Procryptocerus scabriusculus*, *Pseudomyrmex ejectus*, *Pseudomyrmex elongatus*, *Pseudomyrmex filiformis*, *Pseudomyrmex PSW-53*, and *Pseudomyrmex simplex*. We selected a priori to use the 10 most common species. We intended to replicate trials for each pair (out of a total of 45 two-species pairs) at least ten times. However, low encounter rates for some species, and for pairs of species within two days of one another precluded obtaining ideal sample sizes. We replicated trials for each species pair on average 5.73 times; four species pairs were replicated once, nine species pairs were replicated twice, and 31 species pairs were replicated three or more times. Only one species pair (*M. mexicana* and *P. filiformis*) was not tested. We conducted 42 trials in 2007, 105 trials in 2009, 82 trials in 2011, and 30 trials in 2012 for a total of 259 trials.

### Dominance interactions

We used our ant dataset to infer a dominance hierarchy by simulating interactions among individuals to estimate level of uncertainty and steepness in the hierarchy. All simulations were conducted in R version 3.3.3 (R Core Development Team 2017). We used the R package “animDom” version 0.1.2 to infer dominance hierarchies using the randomized Elo-rating method (Sánchez-Tójar et al. 2017). The R package ‘statnet’ was used to test triangle transitivity measures (Handcock et al. 2008).

We subsampled the observed data to determine if the population had been adequately sampled to infer reliable dominance hierarchies. We calculated the ratio of interactions to individuals to determine sampling effort. An average sampling effort ranging from 10-20 interactions is sufficient to infer hierarchies in empirical networks (McDonald and Shizuka 2013). Furthermore, we compared the proportion of observed dyads to the expected proportion of dyads with the probability of interactions of equal group size following a Poisson distribution (Sánchez-Tójar et al. 2017). We estimated the dominance hierarchy using the random Elo-rating method in order to track ranking dynamics over time. We converted the observation data and randomized the order in which sequences of interactions occurred (n = 1000) such that different individual Elo-ratings were calculated each time to obtain mean rankings (Neumann et al. 2011; Strandburg-Peshkin et al. 2015). We estimated uncertainty in the hierarchy by splitting our dataset into two halves and estimated whether the hierarchy in one half of the matrix correlated with the hierarchy of the other half in the matrix (Sánchez-Tójar et al. 2017).

In addition to examining the role of individual ant attributes and levels of uncertainty in dominance hierarchies, we were interested in assessing how the organization of linear dominance hierarchies emerged at higher levels. While individual attributes of species serve as a good predictor of dominance hierarchies among dyadic pairs, the pattern becomes less clear when seeking to explain linear dominance hierarchies at higher levels of species interactions (Chase and Seitz 2011). Therefore, we examined the formation of dominance hierarchies using motif analysis to identify network structures composed of transitive and cyclical triads (Faust 2007). Motif analysis is commonly used in social network analysis to detect emergent properties of the network structure as an explanation for dominance hierarchies by comparing the relative frequencies of motifs in the observed network to the expected value for the null hypothesis of a random network (Holland and Leinhardt 1972; Faust 2007). We carried out motif analysis with customized randomization procedures (McDonald and Shizuka 2013) to compare the structure of our network model against random network graphs. Species interaction data were represented as a directional outcome matrix. The nodes in the network represent individual ant species and the one-way directional arrows of the edges represent dominant-subordinate relationships. In the random networks, we maintained the same number of nodes and edges as in the observed network, but the directionality and placement of edges were generated randomly. Using the adjacency matrix, we calculated the triad census (McDonald & Shizuka 2012). The triad census allows us to examine directed species interactions (Pinter-Wollman *et al.* 2011). We used the seven possible triad configurations fully composed of three nodes that either have asymmetric or mutual edges (Holland and Leinhardt 1972). We used the network analysis packages ‘statnet’ (Handcock et al. 2008) and ‘Igraph’ (Csardi and Nepusz 2006) in R (R Core Development Team 2017) to calculate the frequencies of triad configurations (total triads = 220) and to compare the observed (N = 10) to the random network graphs (N = 10).

To test for statistical significance between the observed and randomly generated networks, we computed the triangle transitivity (*t*_*tri*_). Although the triad census consists of 8 different triangle configurations, we focus our attention on the relative frequencies of transitive triangles. The proportion of transitive triangles P(t) in a dominance network is given by:

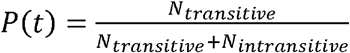

In this case, the expected probability of a transitive triangle in a random network is P(t)=0.75. Using our expected value, we use a scaled index *T*_*tri*_ ranging from 0 as the random expectation to 1 where all triangles are transitive in the network. In random networks the expected frequency of transitive triangles is 0.75. For each empirical network, we simulate 1000 random graphs and calculate the *T*_*tri*_ each time. The P-value represents the number of times the randomized *T*_*tri*_ is greater than the *T*_*tri*_ value of the observed network (Shizuka and McDonald 2012).

## Results

### ‘Real estate’ experiments

In all 258 of 259 trials, there was a clear winner of the ‘real estate’ battle after 24 hours, meaning that one of the two species had occupied the artificial nest. From examining the wins and losses, a clear hierarchy emerged, with some species winning the vast majority (>70%) of trials in which they were involved, and other species winning few trials (<30%). The ranking shows that the twig-nesting species *Myrmelachista mexicana* is the highest ranked species, while *Pseudomyrmex ejectus* is the lowest ranked species in the hierarchy (Table 1). The one trial that did not result in a winner was a trial involving *P. elongatus* and *P. ejectus.*

**Table 1.** Estimation of dominance hierarchy using Elo-rating method. Using 10 ant species in the competition experiments, we recorded a total of 258 interactions. The ratio of interactions to species indicates that the sampling effort of 25.8 falls within the minimum 10-20 range. The ranking shows that the twig-nesting species *Myrmelachista mexicana* is the highest ranked species, while *Pseudomyrmex ejectus* is the lowest ranked species in the hierarchy. Species are as follows: Myrm (*Myrmelachista mexicana*), Ps53 (*Pseudomyrmex* PSW-53), Neso (*Nesomyrmex echinatinodis*), Ca.ab. (*Camponotus abditus*), Ca.n. (*Camponotus colobopsis*), Psfili (*Pseudomyrmex filiformis*), Pssimp (*Pseudomyrmex simplex*), Procryp (*Procryptocerus scabriusculus*), Pselong (*Pseudomyrmex elongatus*), Psej (*Pseudomyrmex ejectus*).

| Table 1. Estimation of dominance hierarchy using Elo-rating method |  |  |  |  |  |  |  |  |  |  |
| --- | --- | --- | --- | --- | --- | --- | --- | --- | --- | --- |
| Species | Myrmelachista mexicana | Pseudomyrmex (PSW-53) | Nesomyrmex echinatinodis | Camponotus abditus | Camponotus colobopsis | Pseudomyrmex filiformis | Pseudomyrmex simplex | Procryptocerus scabriusculus | Pseudomyrmex elongatus | Pseudomyrmex ejectus |
| Rankings | 1.105 | 2.290 | 2.792 | 4.445 | 5.231 | 5.375 | 7.579 | 8.081 | 8.154 | 9.948 |

### Dyadic interactions: Estimating Uncertainty in Dominance Hierarchy

The total number of interactions among the ten species was 258 interactions. The ratio of interaction to individuals (25.8) shows an adequate sampling effort beyond the 10-20 recommended range (Sánchez-Tójar et al. 2017). Using the randomized Elo-rating method, we found that the shape of the hierarchy was intermediate in its steepness showing that rank in the hierarchy largely predicts the probability of winning an interaction (Fig 1). We quantify this by using the Elo-rating repeatability and found a steepness of 0.938.

**Figure 1.**
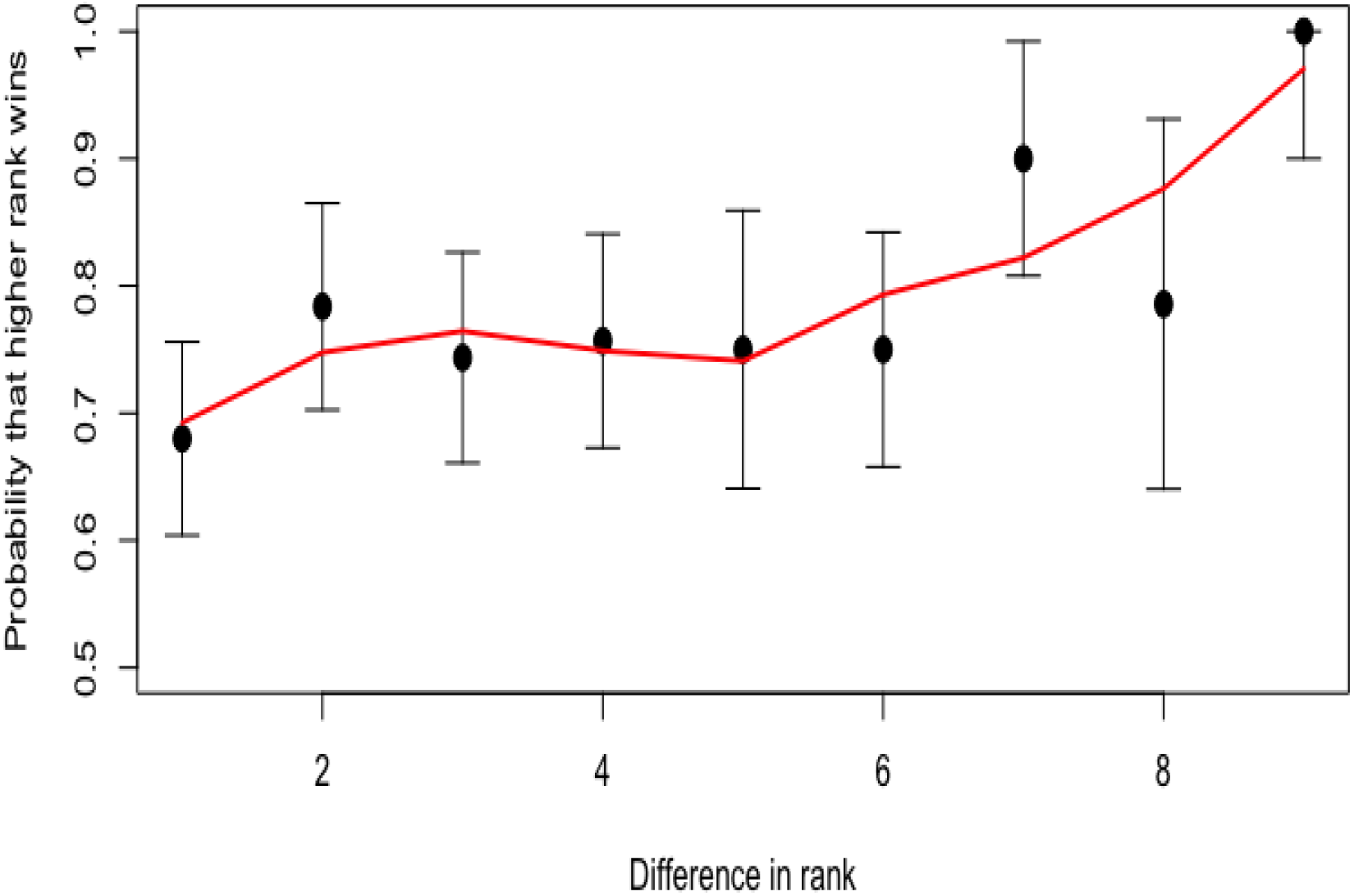
The probability of a higher ranked individual winning. The shape of the hierarchy indicates that the rank is intermediate. We quantified the uncertainty/steepness of the hierarchy based on Elo-rating repeatability which is independent of group size and the ratio of interactions to individuals (Sanchez-Tojar et al. 2017). Based on the Elo-rating, we find that the value obtained is 0.938 which corroborates our qualitative results showing that the hierarchy is intermediate. Thus, rank in this network is a relatively good predictor that a higher ranked individual is more like to win from lower-ranked individuals even though that is not always the case.

We further estimated the uncertainty in the hierarchy by splitting the database into two, and estimating whether hierarchy from one half resembles the hierarchy estimated from the other half. We find that the degree of uncertainty/steepness in the hierarchy is intermediate (mean=0.76, 2.5 % and 97.5% quantile = (0.50,0.94)).

### Triad census analysis

The triad census analysis of the triad distribution showed that the observed network has a significant excess of transitive triads (N=76) followed by a significant deficit of cyclical triads (N=8). Triad types that are positive (i.e. non-overlapping at 0) occurred in excess in the observed network, while triad types that are negative showed a deficit in the observed network as compared to the random null network (Fig 2).

**Figure 2.**
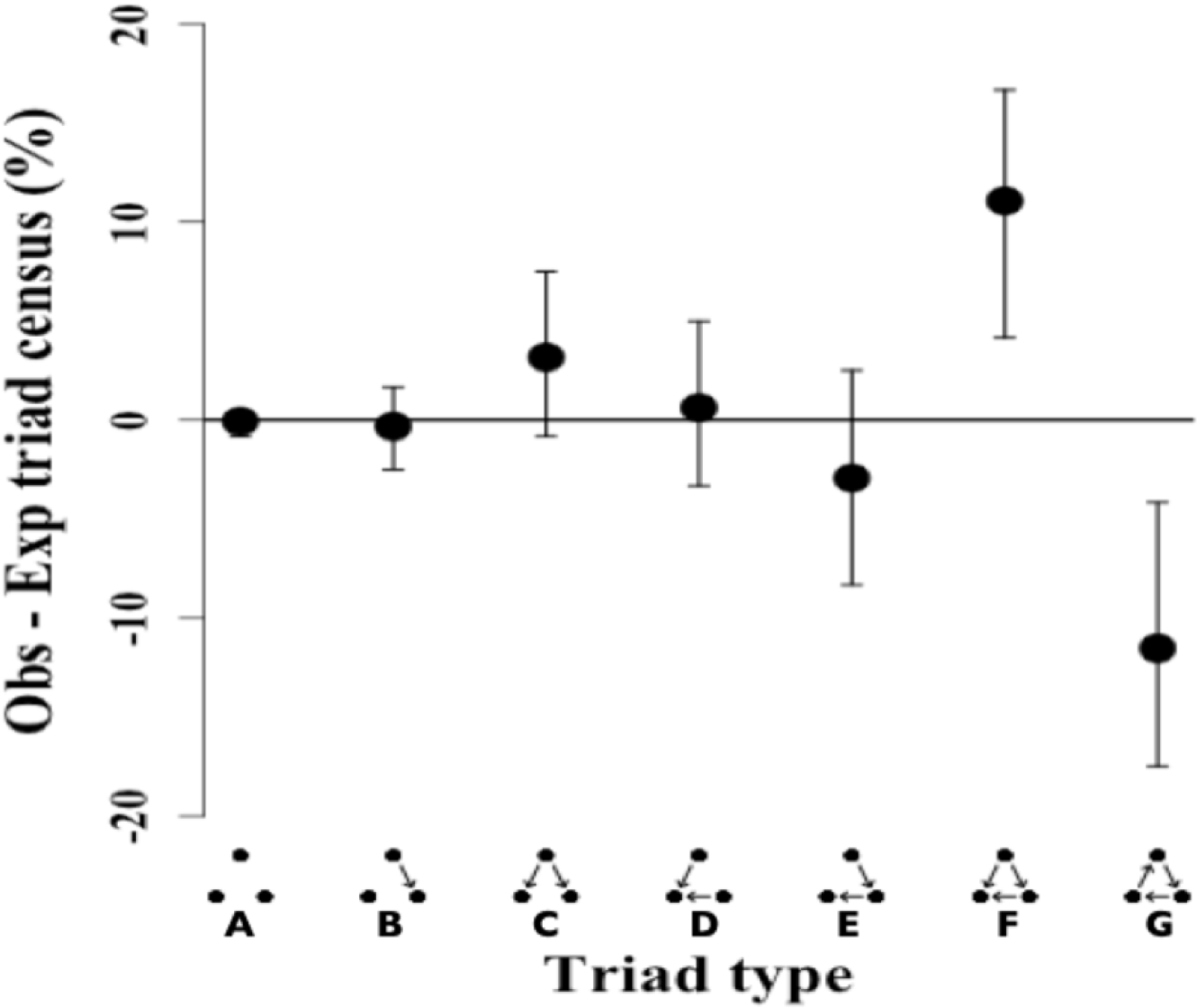
Triad census of twig-nesting arboreal ants. We determined the orderliness of hierarchy by estimating the transitivity of interactions. The y-axis represents the mean difference between the observed (ten ant species network) and expected (10,000 random networks) percentage of the triad subtypes (shown on the x-axis) and error bars show 95% confidence intervals. The twig-nesting ant data show a significant excess of transitive triads (transitive=0.66, p-value=0.002) and a significant deficit of cyclical triads. All the other triad sub-types found were not significantly different from the expected random network. The following symbols define seven possible triad types: A= Null, B=Single-edge, C=Double-dominance, D=Double-subordinate, E=Pass-along, F=Transitive, G=Cycle.

The remaining five triads in the network did not show any significant differences in the mean triad percentage rates between the observed and expected network. While the data showed a clear excess of transitive triangles (34.55 %) and deficit for cyclical triangles (3.6%), the distribution for pass-along triads shows a less typical pattern with the 95% confidence intervals crossing the zero line but the mean percentage still showing a deficit.

## Discussion

Tropical arboreal twig-nesting ants formed a linear dominance hierarchy in our study system. We found that *M. mexicana* was ranked the highest, using Elo-rating method, while *P. ejectus* species ranked the lowest in the hierarchy. We further estimated the level of uncertainty associated with the hierarchy and found an intermediate level of steepness in dominance suggesting a relatively stable hierarchy. The randomized Elo-rating repeatability was 0.93 and remained very stable independently of the ratio of interactions to individuals. To verify that this pattern was not due to a lack of sampling effort, we correlated the two halves of the interaction dataset and found that our sampling effort was 25.8, falling above the minimal recommended range of 10-20 species. Overall, we find that the probability of a higher ranked individual winning a contest against a lower ranked is relatively high, which corroborates our qualitative finding of the hierarchy steepness. Motif analysis of the network further revealed a significant excess of transitive interactions. Thus, the degree of orderliness in the hierarchy is maintained at higher levels of interactions, or in other words, dominance is relatively linear, and transitive, indicating that there is a strong, structured hierarchy in nest-site competition for arboreal twig-nesting ants in coffee agroecosystems.

A question stemming from these results is what effect this might have on structuring the distribution and relative abundance patterns of ants under natural conditions. In Mediterranean ecosystems, dominant and subordinate ants are partitioned on the basis of their life-history traits (Arnan et al. 2012). Dominant ant species had more abundant colonies and displayed increased defense for resources in contrast to subordinate ant species. Meanwhile, subordinate ants exemplified greater tolerance to higher temperatures (Cros et al. 1997; Cerdá et al. 1998*b*). The outcome of interspecific interactions within the dominance hierarchy was contingent on environmental conditions (Arnan et al. 2012). In a temperate forest ecosystem of North Carolina, dominance was context dependent (Stuble et al. 2017). Rankings on the basis of bait monopolization revealed that dominance correlated positively with relative abundances since the most abundant species were ranked higher in the dominance hierarchy. In contrast, rankings based on aggressive encounters did not correlate with abundance. In some habitats, dominance patterns are largely determined by the time of day that foraging occurs (Cerdá et al. 1998*b*; Bestelmeyer 2000). In the North Carolina system, the most abundant ant species *Aphaenogaster rudis* was most active during the morning hours, whereas the cold-tolerant *Prenolepis imparis* species was dominant during the night hours (Stuble et al. 2017). The ranking of species also depends on the size of resources. In an assemblage of woodland ants, smaller-sized ants were more efficient at acquiring and transporting fixed resources. Bigger sized solitary ants excelled at retrieving smaller food that were mobile during competitive interactions. However, the introduction of phorid parasitoids in this system reduced the transitive hierarchy facilitating the coexistence of subdominant ants (LeBrun 2005; Lebrun and Feener 2007). In our study on competition for nesting sites in the lab, we were able to used fixed resources and to a certain degree control variation in colony size.

Regardless of the ecological factors driving dominance hierarchies under natural conditions, it’s important to note that ranking methods vary considerably among studies (Stuble et al. 2013). Traditionally, field studies have quantified dominance relationships on the basis of proportion of contests won. Other studies have use more sophisticated methods to account for competitive reversals (Vries 1998) or have updated rankings based on relative wins and losses during contests (Colley 2002). In this study, we used the Elo-rating by randomizing the sequence of interactions and calculating the mean of individual Elo-ratings (Sánchez-Tójar et al. 2017). By calculating the repeatability of the individual Elo-rankings, we estimated the uncertainty in the rankings by obtaining confidence intervals for each species. This uncertainty measure has allowed us to detect an intermediate dominance hierarchy. By further considering ant communities as networks of interacting species, we have found that twig-nesting ants are overwhelming self-organized into a transitive dominance hierarchy.

More broadly, a high proportion of stable transitive relations have been observed in other animal systems for both dyadic and triadic level interactions (McDonald and Shizuka 2013; Shizuka and McDonald 2015). For example, dominance contest over free-ranging African elephants in Kenya showed that between-group dominance structure is highly transitive (Wittemyer and Getz 2007). Despite the wide geographical distribution of resources and infrequent contests among elephants, the potential cost for conflict were sufficiently high resulting in strong winner and loser effects. Among wild chimpanzees, both male and females formed dominance hierarchies. However, for high ranking female chimpanzees, dominances was associated with reproductive fitness due to greater access to food resources (Wittig and Boesch 2003). Miller et al. 2017 partnered with citizen scientists to examine social hierarchies among bird species at the continental scale. They used extensive catalogs of interspecific interactions at bird feeders across North America and observed that, across the continent, birds at feeders formed a linear dominance hierarchy encompassing an ecologically diverse range of species. Social dominance within a hierarchy was found to be strongly associated with body size with higher ranked species having preferential access to food resources at feeders.

While our study shows that nest site competition is important for structuring twig-nesting ant communities, dominance hierarchies are often context-dependent and ranking of the same species varies across geographical regions or disturbance regimes (Andersen 1997; Feener *et al.* 2008, Sensenig *et al.* 2017). Previous research involving ant competition for variable resources in temperate ecosystems showed that intransitive competition at local spatial scales mediates ant coexistence (Sanders and Gordon 2003). Microclimatic factors also disrupt dominance hierarchies (Cerdá et al. 1998*b*). For instance, environmental variation in agricultural coffee systems is likely to influence dominance hierarchies (Philpott and Foster 2005*a*; Perfecto and Vandermeer 2011). Likewise, occurrence of fire can disrupt dominance hierarchies in specialist ants in *Acacia* trees resulting in increased abundance of subordinate ants (Sensenig et al. 2017) Further, top down processes such as predation and parasitism are likely to mediate twig-nesting ant competition in natural communities (Philpott et al. 2004; Feener et al. 2008*b*; Hsieh and Perfecto 2012). In addition, competition and disturbance from ground- and carton-nesting ants may influence the colonization and community composition of arboreal twig-nesting ants (Philpott et al. 2004*a*; Ennis and Philpott 2017). Therefore, more comparative research is needed to examine how geographical differences or disturbance affects the hierarchy and ultimately the distribution and relative abundance of different arboreal, twig-nesting ant species.

A myriad of other factors may drive the distribution and relative abundance of arboreal ants (Yamaguchi 1992*a*; Palmer et al. 2000*a*). Variation in life-history trade-offs can influence dominance patterns. For example, competition-colonization trade-offs have been identified between competitive colonies expanding unto nearby trees and foundress queens establishing new nest sites (Stanton et al. 2002*a*). Twig-ant communities are strongly influenced by canopy structure and habitat complexity (Philpott et al. 2018). Tree size correlates positively with ant abundance (Yusah and Foster 2006), species richness (Klimes et al. 2015), and composition (Dejean et al. 2008). Canopy connectivity, in turn, impacts local species coexistence as lower connectivity decreases species richness (Powell et al. 2011). Canopy connections serve as pathways by which arboreal ants access tree resources. Limited access to nesting cavities hampers growth and reproduction of arboreal ants (Philpott and Foster 2005). Differences in nest entrances can affect both the abundance and richness of arboreal ant species that are competing for cavities resources (Jiménez-Soto and Philpott 2015). A study examining the effects of twig diversity on litter-nesting ant species found that twigs derived from different trees species harbored greater ant diversity as compared to twigs obtained from a single tree species (Armbrecht et al. 2004). In some ant species belonging to the genus *Cephalotes,* ant size and nest entrance size impacted survival and colony fitness (Powell 2009). Although our artificial twigs featured standardized cavity entrances, variations in ant sizes and cavity entrance size in natural conditions will be important in determining arboreal ant dominance. Thus, future studies should consider variation in nest entrance size to guide our understanding in twig dominance hierarchies.

Based on our findings, we find that tropical twig-nesting ants competing for nesting resources are arranged in a dominance hierarchy. We find that the arboreal ants *Myrmelachista mexicana* was the highest ranked species in the hierarchy, while *Pseudomyrmex ejectus* was ranked as the lowest in the hierarchy. To more reliably infer a hierarchy in our dataset, we accounted for uncertainty in the hierarchy by estimating the probability that a higher ranked defeats a subordinate species. We found that an intermediate steepness level characterized the hierarchy after corroborating an adequate sampling effort. Moving beyond simple pair-wise interactions, we used motif analysis to infer higher order interactions. Transitive interactions were significantly over-represented in the network which further illustrates that twig-nesting ants are organized in a linear hierarchy. While we find that twig-nesting ants form a dominance hierarchy in this tropical agricultural system, it’s likely to expect variation in domination patterns across ecosystems and habitats. Subsequent studies should link dominance patterns with relative abundance patterns in the field in order to assess if particular species traits are important in structuring local communities. While competitive outcomes in our experiment are static (winner and loser), dominance hierarchies exhibit considerable variation and field studies should therefore include spatial and temporal variation. Dominance hierarchy studies are typically designed to assess antagonistic interactions, but less focus has been placed on collecting data with neutral interactions. Difference in food preference and temporal foraging patterns suggest that species don’t necessarily interact in an antagonistic fashion. Therefore, more studies noting neutral interactions will shed greater light on the prevalence of dominance hierarchies under natural conditions.

## Acknowledgements

The following people assisted with field and lab data collection: G. Domínguez Martínez, U. Perez Vasquez, G. Lopez Bautista, F. Sanchez-Lopez, D. Lopez, P. Bichier, B. Chilel, A. De la Mora, D. Gonthier, G. Livingston, K. Mathis, K. Ennis, E. Jimenez-Soto, J. Vandermeer, I. Perfecto, D. Jackson, H. Hsieh, and A. Iverson. J. Rojas and E. Chamé Vasquez of El Colegio de la Frontera Sur (ECOSUR) provided logistical support. Permission for arthropod collection was granted by SEMARNAT (Secretaria de Medio Ambiente y Recursos Naturales). We thank Finca Irlanda and Don Walter Peters for access to the farm and housing for field research. We also wish to thank the participants of the NIMBios workshop on “Animal Social Networks”. Funding was provided by NSF DEB-1262086 to SP, Rackham Merritt Fellowship to SY, and EEB Block Grant to SY.

## Data Accessibility

Twig-ant competition data and scripts to calculate dominance rankings are available via github https://github.com/shenii/twignesting_dominance_dataset

